# High-quality *de novo* genome assembly of the *Dekkera bruxellensis* UMY321 yeast isolate using Nanopore MinION sequencing

**DOI:** 10.1101/151167

**Authors:** Téo Fournier, Jean-Sébastien Gounot, Kelle Freel, Corinne Cruaud, Arnaud Lemainque, Jean-Marc Aury, Patrick Wincker, Joseph Schacherer, Anne Friedrich

## Abstract

Genetic variation in natural populations represents the raw material for phenotypic diversity. Species-wide characterization of genetic variants is crucial to have a deeper insight into the genotype-phenotype relationship. With the advent of new sequencing strategies and more recently the release of long-read sequencing platforms, it is now possible to explore the genetic diversity of any non-model organisms, representing a fundamental resource for biological research. In the frame of population genomic surveys, a first step is evidently to obtain the complete sequence and high quality assembly of a reference genome. Here, we completely sequenced and assembled a reference genome of the non-conventional *Dekkera bruxellensis* yeast. While this species is a major cause of wine spoilage, it paradoxically contributes to the specific flavor profile of some Belgium beers. In addition, an extreme karyotype variability is observed across natural isolates, highlighting that *D. bruxellensis* genome is very dynamic. The whole genome of the *D. bruxellensis* UMY321 isolate was sequenced using a combination of Nanopore long-read and Illumina short-read sequencing data. We generated the most complete and contiguous *de novo* assembly of *D. bruxellensis* to date and obtained a first glimpse into the genomic variability within this species by comparing the sequences of several isolates. This genome sequence is therefore of high value for population genomic surveys and represents a reference to study genome dynamic in this yeast species.

## Introduction

Knowledge in biology has been greatly improved by exploring a large diversity of species as well as evolutionary contexts. No single species is representative of the evolution of either an entire phylum or a whole genus. Exploration of the genetic diversity of non-model species is essential to have a better insight into the variation of the population history, recombination, selection, mutation, and the genotype-phenotype relationship. In this context, the Saccharomycotina subphylum (budding yeasts), which includes the baker’s yeast *Saccharomyces cerevisiae*, represents an ideal group of non-model organisms for population genomic studies (Peter and Schacherer 2016).

Recent years have seen a burst of population genomic surveys focusing on various non-conventional yeasts associated with different objectives. This has a bearing on several aspects of evolutionary biology. Analysis of resequencing data of a large sample of isolates from the same species has been focused on yeast model organisms such as *Saccharomyces cerevisiae* (Schacherer et al. 2009, Liti et al. 2009, Skelly et al. 2013, Bergstrom et al. 2014, Strope et al. 2015, Almeida et al. 2015, Zu et al. 2016, Gallone et al. 2016, Gonçalves et al. 2016) and the fission yeast *Schizosaccharomyces pombe* (Fawcett et al. 2014, Jeffares et al. 2015), as well as on non-model yeast species: *Saccharomyces paradoxus* (Leducq et al. 2016), *Saccharomyces uvarum* (Almeida et al. 2014), *Candida albicans* (Hirakawa et al. 2015, Ford et al. 2015) and *Lachancea kluyveri* (Friedrich et al. 2015, Brion et al. 2015, 2016). Altogether, these data and analysis enhanced our knowledge about the evolutionary history of species (Almeida et al. 2014), the forces involved in genome evolution (Friedrich et al. 2015), and the genetic basis of the phenotypic diversity (Ford et al. 2015).

Among the Saccharomycotina, *Dekkera bruxellensis* is a yeast species associated with human fermentation processes well known as a major cause of wine spoilage but also as an essential contributor to Belgium lambic and gueuze beer fermentation (Schifferdecker et al. 2014, Masneuf-Pomarede et al. 2015). In addition to its industrial properties, this species is of interest at the evolutionary level. Natural isolates show different ploidy levels (Borneman et al. 2014, Curtin and Pretorius 2014), and extensive chromosomal rearrangements, which were observed through electrophoretic karyotypes (Hellborg and Piškur 2009). These observations indicate a rapid evolution at the intraspecific level. Recent findings suggest that the ploidy level could be linked to the substrate of origin of the strain and related to adaptive processes linked to specific environments (Albertin et al. 2014). Consequently, a genome-wide polymorphism survey based on a representative set of *D. bruxellensis* individuals would be of interest. The exploration of SNPs (Single Nucleotide Polymorphisms), small indels, but also structural variants such as large indels, inversions and translocations at the species level would help provide insight into the forces that shape genomic architecture and evolution. However, to conduct a population genomic survey, the availability of a high-quality reference sequence for the species at the level of completeness that would cover the majority of the genomic variation but also at the contiguity level to efficiently detect structural variants, is a pre-requisite.

To date, population genomic studies have mostly been performed on species for which chromosomal-scale genome assemblies were available, however, this necessary high-quality assembly was unfortunately not yet available for the *D. bruxellensis* species. Here, we present the de novo sequence and high-quality genome assembly of the UMY321 *D. bruxellensis* isolate with a combination of long Oxford Nanopore and short Illumina reads. By aligning the short-read sequencing data from a total of eight sequenced natural isolates on the generated assembly, as well as other previously available assemblies (Piskur et al. 2012, Curtin et al. 2012, Borneman et al. 2014, Crauwels et al. 2014, Olsen et al., 2015), we tested the capacity of our assembly to be used as a reference assembly for future population genomic studies of this non-model species. The results showed that we generated the most complete and contiguous *de novo* assembly of *D. bruxellensis*, necessary to explore the intraspecific genetic diversity of this unique and economically relevant species.

## Materials and Method

### Yeast strains and DNA preparation

We selected three *D. bruxellensis* diploid isolates from various ecological and geographical origins (Table 1). The UMY321 isolate was chosen for the generation of a high-quality assembly and was therefore subjected to Oxford Nanopore and Illumina sequencing. The two other isolates, UMY315 and 133, were only subjected to Illumina sequencing for comparative analysis purposes.

**Table 1:** Description of the *D. bruxellensis* isolates used in this study

| Strain | Ploidy | Ecological origin | Geographical origin | Reference |
| --- | --- | --- | --- | --- |
| AWRI1499 | 3n | wine | Australia | Curtin et al., 2012 |
| AWRI1608 | 3n | wine | Australia | Borneman et al., 2014 |
| AWRI1613 | 2n | wine | Australia | Borneman et al., 2014 |
| CBS11270 | 2n | industrial ethanol | Sweden | Olsen et al., 2015 |
| CBS2499 | 2n | wine | France | Piskur et al., 2012 |
| ST05_12_22 | 2n | lambic beer | Belgium | Crauwels et al., 2014 |
| UMY315 | 2n | must | Italy | this study |
| UMY321 | 2n | red wine | Italy | this study |
| 133 | 2n | merlot wine | South Africa | this study |

Yeast cell cultures were grown overnight at 30°C in 20 mL of YPD medium to early stationary phase before cells were harvested by centrifugation. Total genomic DNA was than extracted using the QIAGEN Genomic-tip 100/G according to the manufacturer’s instructions.

### Flow cytometry

Samples were prepared for DNA content analysis using flow cytometry. Cells were grown in YPD medium at 30°C to reach exponential phase. They were then pelleted and washed with 1 ml of water. In order to fix the cell, pellet was resuspended in 1ml of 70 % ethanol. After centrifugation, supernatant was discarded and cells were resuspended in 1 ml of sodium citrate buffer (tri-sodium citrate 50 mM; pH 7.5). Cells were pelleted once more and resuspended in 1 ml sodium citrate buffer supplemented with 10 μl of RNase A (100 mg/ml) and incubated at 37°C for 2 hours. Samples were then sonicated (Sonics® Vibra-Cell VC750) for 20 seconds with a 20 % amplitude. After sonication, 1 ml of sodium citrate buffer supplemented with 10 μl of propidium iodide (1.6 mg/ml) and left in the dark at 4°C for 12 hours. Once the cells were stained with propidium iodide, cells DNA content was assessed by measuring fluorescence intensity using flow cytometry (Cyflow® Space, Partec).

### MinION library preparation and sequencing

Two μg of genomic DNA was sheared to approximately 8,000 bp with g-TUBE. After clean-up using 1x AMPure XP beads, Nanopore’s 8 kb 2D sequencing libraries were prepared according to the SQK-MAP005-MinION gDNA Sequencing Kit protocol.

The sequencing mix was prepared with 8 μL of the DNA library, water, the Fuel Mix and the Running buffer according to the SQK-MAP005 protocol. The sequencing mix was added to the R7.3 flowcell for a 48 hours run. The flowcell was reloaded one time at 24 hours with an addition of 8 μL of the DNA library.

### Illumina Sequencing

Genomic Illumina sequencing libraries were prepared with a mean insert size of 280 bp and were subjected to paired-end sequencing (2 x 100 bp) on Illumina HiSeq2000 sequencers.

### *de novo* genome assembly

Various sets of the longest MinION 2D reads, which refer to various theoretical genome coverage (10x, 15x, 20x and all the 2D reads) taking 15 Mb as genome size estimate (Table S1) were subjected to four assemblers: ABruijn (v0.3b) (Lin et al. 2016) Canu (v1.1) (Berlin et al. 2015), miniasm (v0.2- r137-dirty) (Li et al. 2016) and SMARTdenovo (https://github.com/ruanjue/smartdenovo). ABruijn and miniasm were run with default parameters, while “genomeSize=13m, minReadLength=2500, mhapSensitivity=high, corMhapSensitivity=high and corOutCoverage=500” was set for Canu and “-c 1 -k 14 -J 2500 -e zmo” for SMARTdenovo. After the assembly step, we polished each set of contigs with Pilon (v1.18) (Walker BJ et al. 2014), using ~100X of Illumina 2 x 100 bp paired-end reads. SSPACE-LongRead (v1.1) (Boetzer and Pirovano 2014) was finally used to scaffold the selected assembly using long-reads information.

### Whole genome comparison

Whole genome comparisons were performed with MUMmer (v3.0) (Kurtz et al. 2004). nucmer was used to align the sequences (with --maxmatch option). The alignments coordinates were extracted to determine the proportion of non-N residues of each assembly that were covered. The delta files were filtered for alignments < 5kb and plots were generated with mummerplot.

### Short reads mapping

Reads were mapped with BWA (V0.7.4) (Li and Durbin 2009) and unmapped reads were estimated with Samtools (v 0.1.19) (Li et al. 2009). GATK (v3.3) (McKenna et al. 2010) was used for local realignment of the reads around indels, SNPs calling, and to add allele balance information in the vcf file.

### Data availability

All sequencing data generated in this study, as well as the UMY321 reference assembly (in FASTA format) have been deposited in the European Nucleotide Archive under the accession number PRJEB21262.

## Results and Discussion

Three *D. bruxellensis* isolates (UMY321, UMY315 and 133) were sequenced in this study (Table 1). These strains were determined to be diploid based on flow cytometry analysis and were all isolated from wine or grape must in Italy or South Africa. The genome of the UMY321 isolate was sequenced using a combination of Nanopore long-read and Illumina short-read sequencing data to obtain a high-quality assembly. By contrast, the UM315 and 133 isolates were only sequenced using a short read strategy. In addition, these genomes were compared to previously genome sequences of 6 other *D. bruxellensis* isolates (Table 1) (Piskur et al. 2012, Curtin et al. 2012, Borneman et al. 2014, Crauwels et al. 2014, Olsen et al., 2015).

### *de novo* genome assembly construction and comparison

For the UMY321 isolate, a total of three MinION Mk1 runs were performed with the R7.3 chemistry using 2D library types with 8 kb mean fragmentation size. A total of 115,559 reads, representing a cumulative size of 1.15 Gb were generated, among which 41,686 2D reads showed an average quality greater than 9 (2D pass reads). We focused on these 2D pass reads, that represent a total of 376.8 Mb, with the longest read being 70,058 bp (mean = 9,033 bp and median = 8,676 bp) (Figure S1). Four subsets of our 2D pass reads (10x, 15x, 20x of the longest 2D pass reads and all of them) (Table S1) were submitted to four assemblers: ABruijn (Lin et al. 2016), Canu (Berlin et al. 2015), miniasm (Li et al. 2016) and SMARTdenovo (https://github.com/ruanjue/smartdenovo). As the MinION sequencing technology is known to be associated to high error rates (~10% for 2D pass reads) (Jain et al. 2016), we polished the assemblies with Pilon (Walker et al. 2014) using 100x of Illumina paired-end reads. The lengths of the constructed assemblies were all in the same order of magnitude and ranged from 11.7 to 13.7 Mb (Table S2).

Using these various datasets and assemblies, the objective was to define the best assembler and the minimal coverage needed. Hence, we computed the standard contiguity metrics for all assemblies to evaluate their quality, which is related to both the assembler and the dataset (Figure 1, Table S2). First, we observed that, considering the results by assembler, the number of scaffolds obtained with the 10x dataset is much higher compared to the other datasets, which suggests that a 10x coverage of MinION reads is too low to obtain a good quality assembly. By assembler, the results obtained for the higher coverages are comparable. Using Canu, the number of scaffolds is much higher and N90 as well as N50 are much lower, producing the less connected assemblies (Figure 1, Table S2). The contiguity metrics associated to the assemblies constructed with SMARTdenovo, Abruijn and miniasm were closely related, and it seemed difficult to select a single best assembly on the sole basis of these measurements, especially since good contiguity metrics are not necessarily associated with assembly completeness.

**Figure 1.**
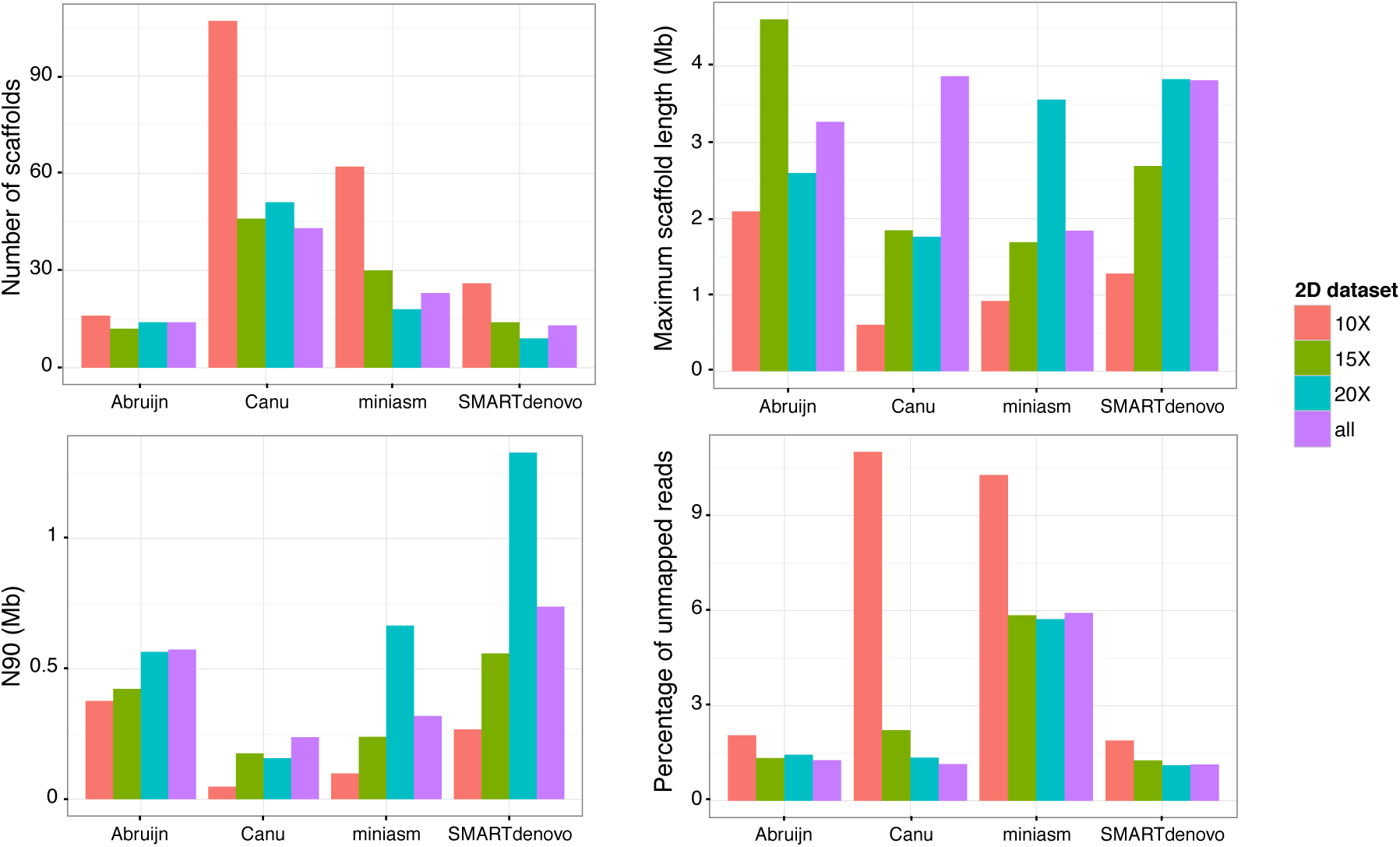
Metrics related to the constructed assemblies, per assembler and dataset.

Considering these results, we decided to map the Illumina PE reads back on the generated assemblies with BWA (Li and Durbin 2009). Among all the assemblies, the proportion of unmapped reads ranged from 1.12 to 11% (Figure 1, Table S2). Surprisingly, the assemblies constructed with miniasm were less complete, as more than 5% of the reads did not map back, compared to less than 1.5% for the Abruijn and SMARTdenovo assemblies.

By comparing standard metrics and the proportion of unmapped reads, the most accurate assembly was obtained with the 20x 2D reads dataset combined with the SMARTdenovo assembler. This assembly is composed of 9 scaffolds, *i.e.* very close to the estimated number of chromosomes, which appears to vary between 4 and 9 among different strains of this species (Hellborg and Piskur, 2009), for a complete assembly size of 12.97 Mb. This latter was then submitted to SSPACE-longreads, which reduced the number of scaffolds to 8 after grouping the 2 smallest ones, based on our long reads information, and a further Pilon run. The final assembly contains 8 scaffolds and shows a cumulative size of 12,965,163 bp (Table 2). We also evaluated the completeness of our assembly at the gene content level by running CEGMA (Parra et al. 2007): 245 out of the 248 most extremely conserved genes in Eukaryotes were detected in our assembly, through 242 complete and 3 partial alignments. Altogether, these results reveal a high level of completeness of our assembly.

**Table 2:**
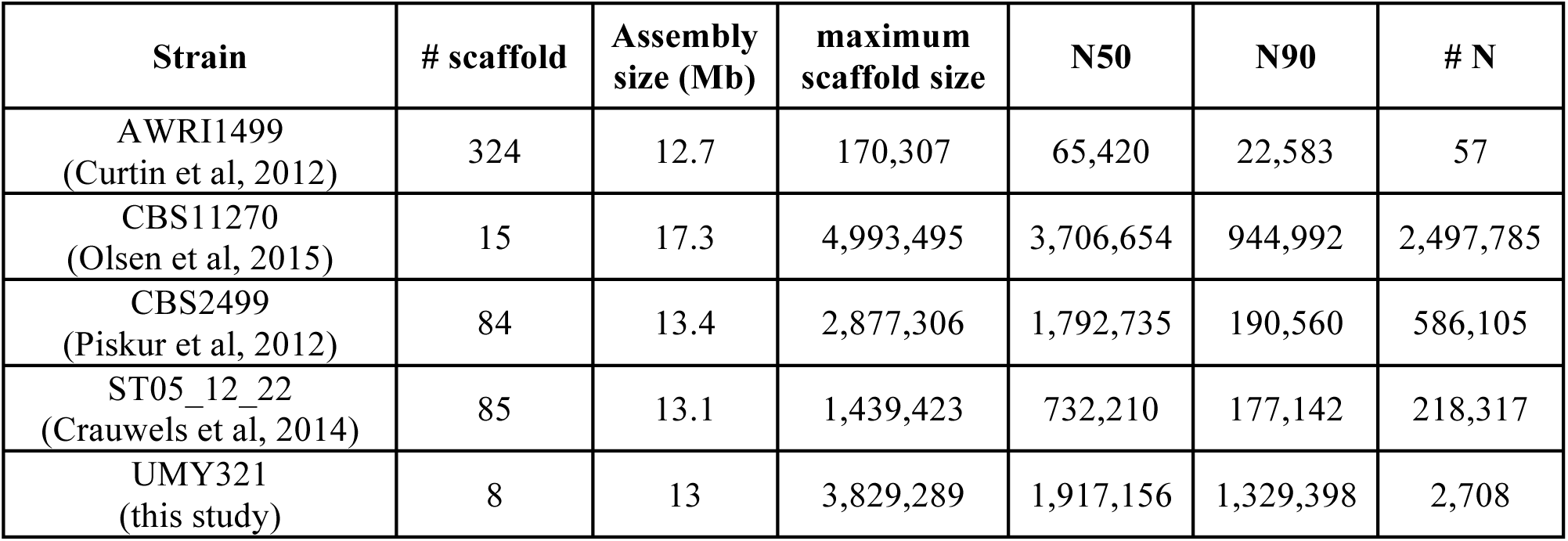
Metrics associated to the *D. bruxellensis* publicly available assemblies

| Strain | # scaffold | Assembly size (Mb) | maximum scaffold size | N50 | N90 | # N |
| --- | --- | --- | --- | --- | --- | --- |
| AWRI1499<br>(Curtin et al, 2012) | 324 | 12.7 | 170,307 | 65,420 | 22,583 | 57 |
| CBS11270<br>(Olsen et al, 2015) | 15 | 17.3 | 4,993,495 | 3,706,654 | 944,992 | 2,497,785 |
| CBS2499<br>(Piskur et al, 2012) | 84 | 13.4 | 2,877,306 | 1,792,735 | 190,560 | 586,105 |
| ST05_12_22<br>(Crauwels et al, 2014) | 85 | 13.1 | 1,439,423 | 732,210 | 177,142 | 218,317 |
| UMY321<br>(this study) | 8 | 13 | 3,829,289 | 1,917,156 | 1,329,398 | 2,708 |

### Comparison with available assemblies of *D. bruxellensis*

To date, several assemblies of the *D. bruxellensis* species have already been released (Piskur et al. 2012, Curtin et al. 2012, Borneman et al. 2014, Crauwels et al. 2014, Olsen et al., 2015). These assemblies are related to isolates from different ecological and geographical origins (Table 1). They were mostly constructed by combining several sequencing methods such as 454, PacBio and Illumina as well as optical mapping in the most recently published one (Olsen et al., 2015).

The assemblies have very variable metrics associated with each of them (Table 2). In terms of contiguity, our assembly and the assembly generated for the CBS11270 isolate are close, and reach a chromosome-scale resolution. However, the CBS11270 assembly is much larger than the others (17.3 Mb *vs* 12.7 to 13.4 Mb), although it does also contain approximately 2.5 Mb of undetermined (N) residues.

By comparing the assembly metrics, we determined that our assembly is closer to that for CBS11270, which was generated by combining PacBio and Illumina sequencing methods as well as optical mapping, and much better than the other 3 available for comparison, which were much more fragmented and comprised at least 84 scaffolds.

A MUMmer comparison of our UMY321 assembly to that of CBS11270 indicates that 91% and 99.6% of the assemblies aligned, respectively, with one another and revealed that the scaffolds are mostly collinear (Figure 2). However, some large repetitive regions can be observed in the CBS11270 assembly, *e.g.* on the chromosome 1, between chromosome 1 and 6, and between chromosome 4 and 5 (Figure 2, Figure S2) that are absent in our assembly, and could explain the size differences between the assemblies (17.3 Mb *vs* 12.97 Mb). Moreover, some synteny breaks can be observed, at the level of scaffolds, specifically between 3 and 4, for example. All the inconstancies between the assemblies could be related either to structural rearrangements between the isolates or to assembly errors, and would require further investigations to reach a conclusion as to their most likely source.

**Figure 2.**
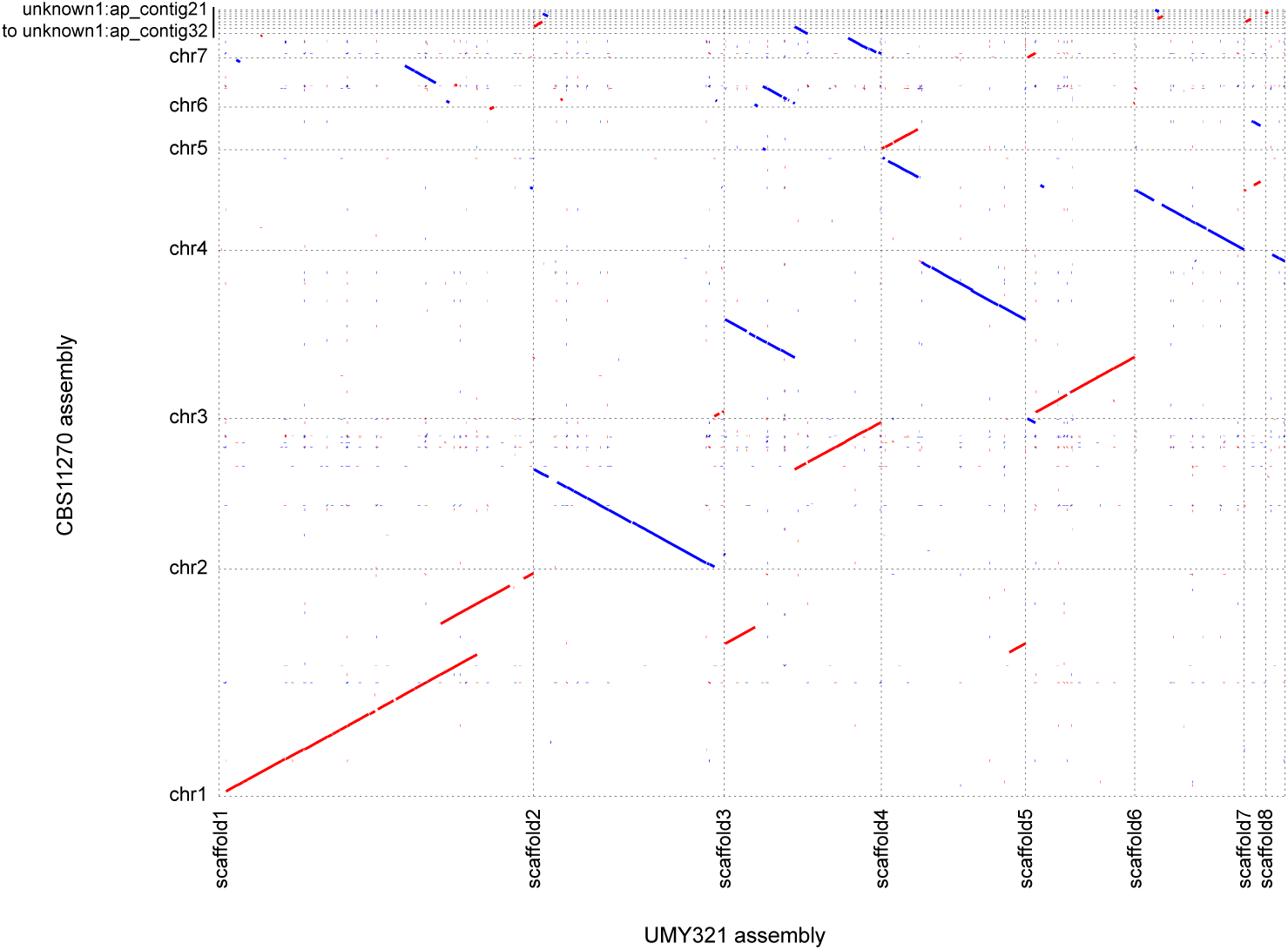
Comparison of the CBS11270 and UMY321 assemblies. The alignments and the plot were generated with the MUMmer software suite.

### Suitability of our assembly for population genomics studies

As previously mentioned, to function as a valuable resource for conducting population genomics studies, a reference genome should combine high-contiguity (for the detection of structural variants) and completeness (for the efficient detection of SNPs and small indels). At the contiguity level, our assembly is close from a chromosomal-scale resolution, which suggests that it would be highly suitable for gross structural rearrangement detection (translocations, inversions, long insertions/deletions).

To test our assembly for the detection of polymorphism along the genome, we further investigated the mapping of the Illumina reads. As previously mentioned, 98.89 % of the UMY321 Illumina reads mapped on our assembly. The read coverage was homogeneous along the scaffolds (Figure 3A), which suggests that the strain is devoid of aneuploidy and segmental duplication, and confirms the lack of large repetitive regions within our assembly.

**Figure 3.**
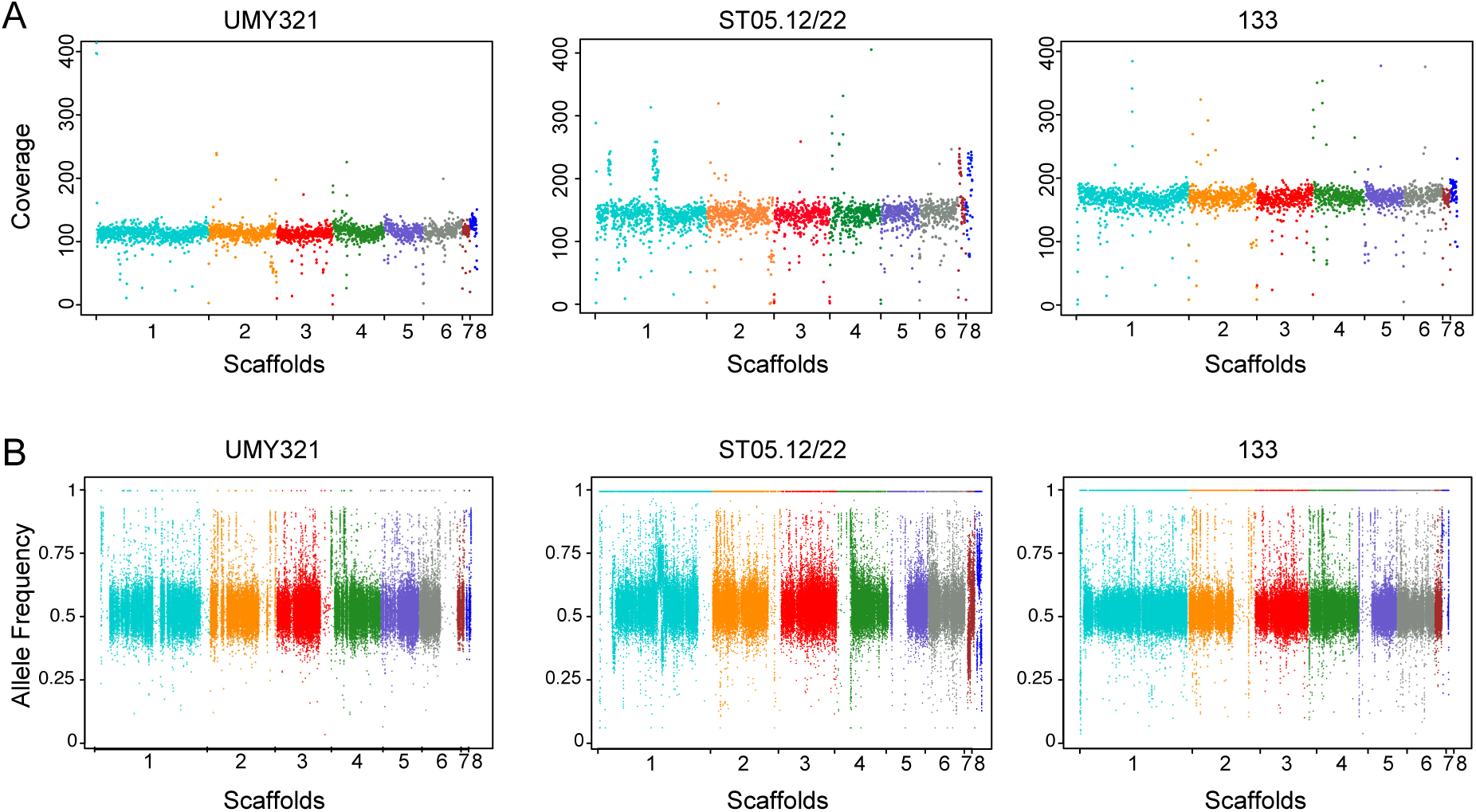
Mapping of the Illumina reads *vs.* the UMY321 reference assembly. (A) Illumina reads coverage along the reference genome (B) Frequency of the reference allele at heterozygous sites along the genome.

A total of 83,006 SNPs were detected with GATK (McKenna et al. 2010), among which 374 were homozygous and 82,632 were heterozygous (Table S3). The 374 homozygous SNPs could be considered as false positives. While not completely negligible, this number is very low and could be related to the high error rate of the MinION technology, which is not completely compensated by using Illumina short reads (Istace et al. 2017).

The UMY321 isolate that we sequenced is diploid, and the detection of these 82,632 heterozygous SNPs revealed that the two genomic copies are not identical and have a high heterozygozity level. These heterozygous positions are mostly evenly distributed all along the genome, with several regions showing loss of heterozygozity (LOH) on scaffolds 1, 2, 3 and 6 (Figure 3B).

Altogether, these results confirmed that our assembly performs well when mapping the reads that were used for its construction. However, to determine if an assembly is relevant in the context of population genomic studies, we also analyzed its performance mapping reads from other isolates. To survey polymorphisms within a species, resequencing projects rely mainly on Illumina sequencing technology, therefore we mapped the short reads related to this species that were publically available as well as from two isolates we sequenced in the context of this project (Table S4) against our assembly and we reported the proportion of unmapped reads. We also aligned these reads against the publically available assemblies to perform a comparative analysis (Table 3). As expected, the UMY321 Illumina PE reads mapped better on our assembly with only 1.11% of unmapped reads. More surprisingly, short-reads generated in the context of the other projects also mapped better on our assembly compared to their related assemblies, and more generally compared to all other assemblies (Figure 4). It is also worth noting that all the reads, including those related to the CBS11270 isolate, mapped less efficiently to the CBS11270 assembly compared to all other assemblies, which suggests that although this assembly is highly contiguous and much larger than the others available, it is less complete.

**Table 3:** Proportion of *D. bruxellensis* unmapped Illumina reads on the available assemblies

|  |  | Assemblies |  |  |  |  |
| --- | --- | --- | --- | --- | --- | --- |
|  |  | UMY321 | CBS11270 | CBS2499 | ST05.12/22 | AWRI1499 |
| Illumina<br>PE reads | UMY321 | 1.11 | 9.95 | 4.13 | 2.12 | 5.29 |
|  | CBS11270 | 4.74 | 12.43 | 5.88 | 3.48 | 9.49 |
|  | CBS2499 | 1.68 | 9.4 | 4.92 | 2.45 | 5.68 |
|  | ST05.12/22 | 1.9 | 10.83 | 7 | 3.78 | 11.97 |
|  | UMY315 | 0.66 | 10.00 | 4.04 | 2.02 | 5.35 |
|  | 133 | 0.82 | 8.82 | 3.11 | 1.57 | 4.44 |
|  | AWRI1608 | 14.87 | 22.65 | 17.42 | 15.39 | 19.91 |
|  | AWRI1613 | 9.69 | 16.89 | 11.04 | 8.98 | 13.38 |

**Figure 4.**
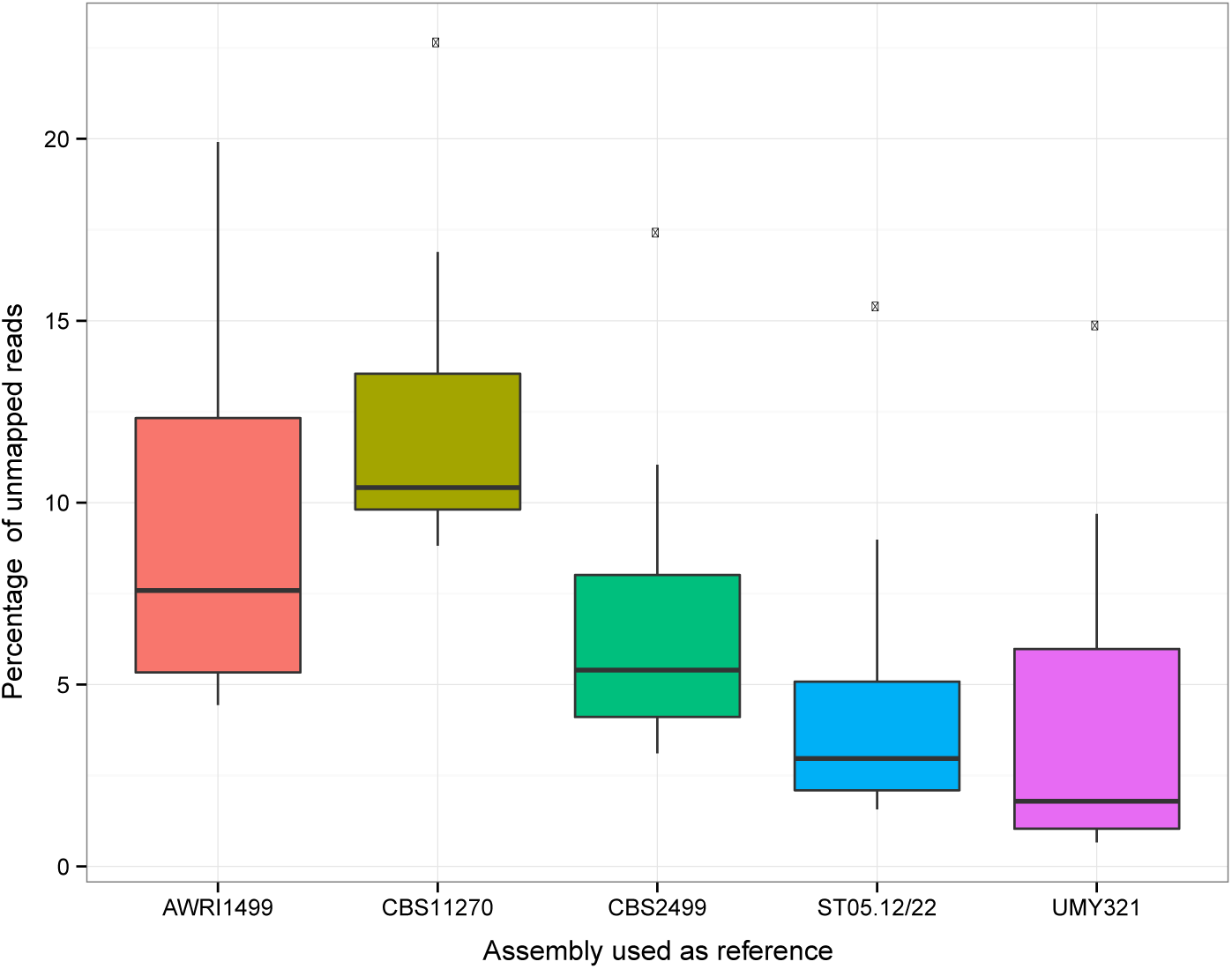
Illumina unmapped reads per assembly. Boxplot of the percentage of unmapped Illumina reads, according to the assembly used for the mapping.

### Insight into the intraspecific genetic variability

Finally, we took advantage of the availability of Illumina reads related to different isolates in order to obtain a first glimpse into the genomic variability within this species, using our UMY321 assembly as a reference. The read coverage along the reference sequence was mostly homogeneous for all isolates, and only few deviations were observed, limited to small genomic regions, which are characteristic of segmental duplications, in the ST05.12/22 isolate (Figure 3A). This suggests that the structural variants within this species are mostly balanced.

Among the 8 studied isolates, one is triploid (AWRI1608) and all the others are diploid (Table 1). A total of 1,268,172 SNPs were detected across these 8 isolates, among which 82% are heterozygous (Table S3). These SNPs are distributed over 500,707 polymorphic positions, with a majority present as singletons (68.8% of the polymorphic sites). However, a significant proportion of this variability is related to the triploid strain AWRI1608. Indeed, when this strain was not included in the analysis, 829,313 SNPs were detected over 188,717 polymorphic positions with only 50,702 singletons (27%). This is in agreement with the proposition that AWRI1608 consists of a slightly heterozygous diploid set of chromosomes with an additional full set of more distantly related chromosomes (Borneman et al. 2014). The phylogenetic relationships between this small sample of isolates based on the whole set of polymorphic positions also reflect the high divergence of this triploid isolate (Figure S3A). Ploidy levels across the genomes were also confirmed by taking advantage of allele frequency at heterozygous positions, which was around 0.5 for diploid isolates and 0.33/0.66 for the AWRI1608 genome (Figure S3B). These heterozygous positions are evenly distributed along the genome, however LOH regions were detected in all the diploid isolates (Figure 3B).

## Conclusion

*D. bruxellensis* is a yeast species of great importance in fermented beverage industries, largely thought of as a contaminant organism (Schifferdecker et al. 2014, Masneuf-Pomarede et al. 2015). This species is also an interesting model to study genome evolution and dynamics as it is characterized by a large genomic plasticity. For these reasons, we sought to generate a high-quality genome assembly and ultimately obtain a suitable reference genome for population genomics. Our analyses show that the *D. bruxellensis* assembly that we generated with a combination of moderate coverage (20x) MinION long-reads in addition to higher coverage (100x) of Illumina reads utilized for sequence polishing purposes, is highly valuable for population genomic studies and outperforms previously publically available sequences. Preliminary comparison among a small set of 9 isolates already highlights the presence of large regions of loss of heterozygosity, which appears to be key factor in the genome evolution and adaptation of a large number of yeast species (Magwene et al. 2011, Ford et al. 2015, Smukowski Heil et al. 2017). To obtain a species-wide view of the genetic variability of *D. bruxellensis*, many more isolates should be surveyed using both short-read as well as long-read sequencing techniques, which will allow for the exploration of the structural variant landscape.

## Acknowledgements

We are grateful to Warren Albertin and Isabelle Masneuf-Pomarede for fruitful discussions and invaluable advices. We thank Jure Piskur and Anna Schifferdecker for generously providing the UMY321, UMY315 and 133 *Dekkera bruxellenis* isolates. We also thank the BioImage platform (IBMP-CNRS, Strasbourg) for their support. This work was supported by France Génomique (ANR-10-INBS-09-08), and the Agence Nationale de la Recherche (ANR-16-CE12-0019 to JS). TF and JSG are supported by a grant from the French “Ministère de l’Enseignement Supérieur et de la Recherche”. JS is a Fellow of the University of Strasbourg Institute for Advanced Study (USIAS) and a member of the Institut Universitaire de France.

## Supplementary figures

**Figure S1.**
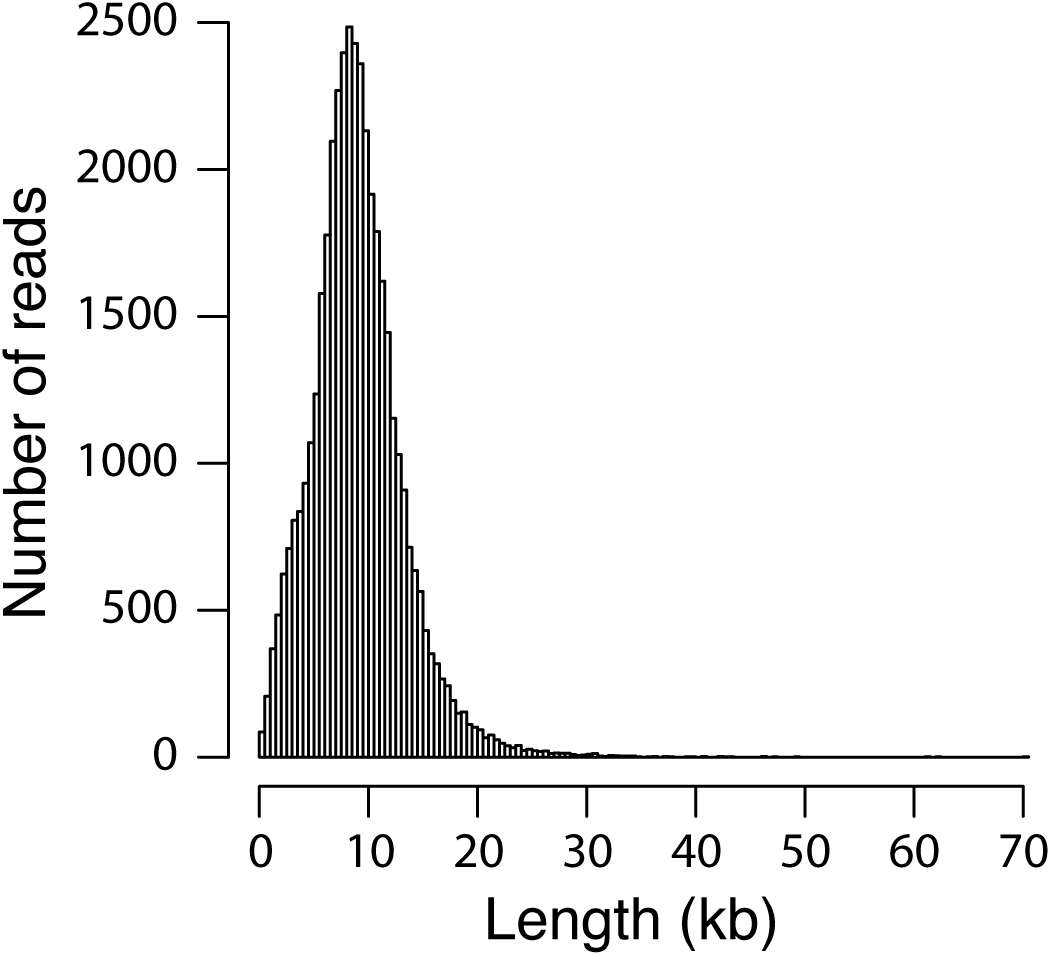
Distribution of the length of the 41,686 2D pass reads.

**Figure S2.**
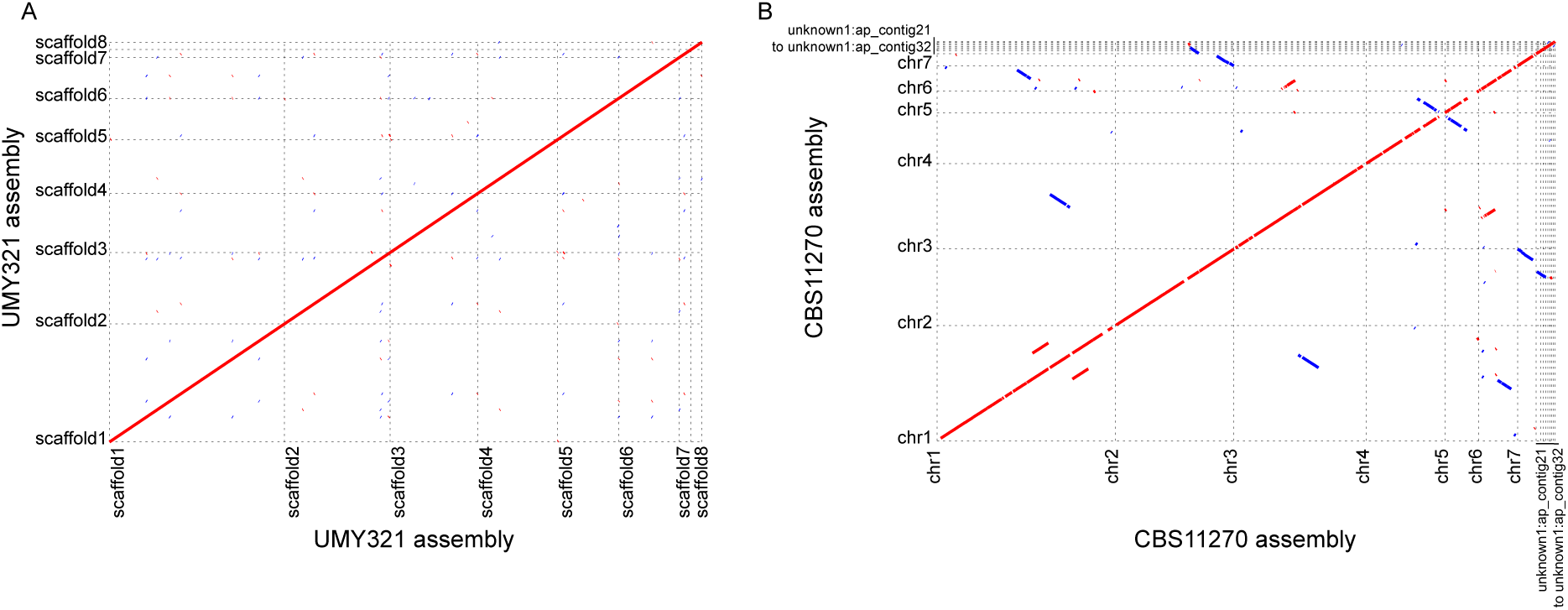
MUMmer-based genomic comparison of assemblies for large repeated regions detection. A. The UMY321 assembly is compared to itself. B. The CBS11270 assembly is compared to itself.

**Figure S3.**
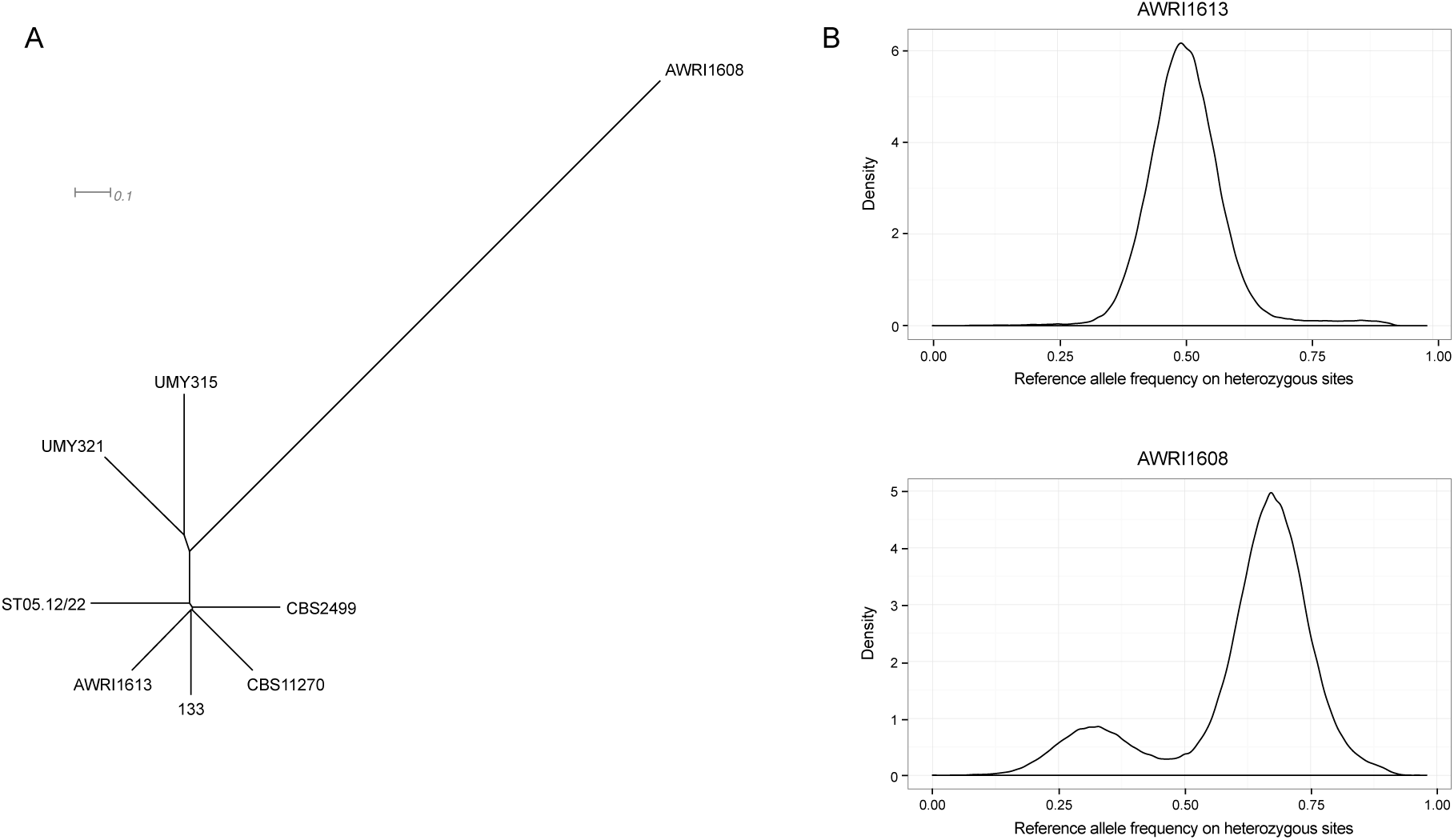
Intraspecific genetic variability. (A) Neighbor-joining tree of *D. bruxellensis* isolates constructed based on 500,707 polymorphic sites. (B) Density of the reference allele frequency for a diploid isolate (AWRI1613) and for a triploid isolate (AWRI1608).

